# An Agent-based model of the gradual emergence of modern linguistic complexity

**DOI:** 10.1101/2020.11.12.380683

**Authors:** Marcel Ruland, Alejandro Andirkó, Iza Romanowska, Cedric Boeckx

## Abstract

A central question in the evolution of human language is whether it emerged as a result of one specific event or from a mosaic-like constellation of different phenomena and their interactions. Three potential processes have been identified by recent research as the potential *primum mobile* for the origins of modern linguistic complexity: *Self-domestication*, characterized by a reduction in reactive aggression and often associated with a gracilization of the face; changes in early brain development manifested by *globularization* of the skull; and *demographic expansion* of *H. sapiens* during the Middle Pleistocene. We developed an agent-based model to investigate how these three factors influence transmission of information within a population. Our model shows that there is an optimal degree of both hostility and mental capacity at which the amount of transmitted information is the largest. It also shows that linguistic communities formed within the population are strongest under circumstances where individuals have high levels of cognitive capacity available for information processing and there is at least a certain degree of hostility present. In contrast, we find no significant effects related to population size.

## Background

By any metric, the modern human mental capacity for language—our ability to construct grammars, accumulate vast vocabularies and flexibly deploy them in relaying meaning—generates complex outputs that go far beyond what other animal communication systems display. For a long time linguists, archaeologists and researchers in adjacent disciplines looked for a single element or event that could have triggered the emergence of the modern human capacity for language (Klein, 2009; Berwick and Chomsky, 2016), taken as the signature of ‘cognitive modernity’. Despite its appeal, this view of a single cause appears to be increasingly implausible (Martins and Boeckx, 2019; De Boer et al., 2020) and a wide range of factors are thought to have shaped our language abilities, ranging from the biological to the cultural and the ecological (Tallerman and Gibson, 2012), more in line with the mosaic nature of the evolutionary trajectory of our species (Scerri et al., 2018).

Understanding the interdependence of multiple factors and revealing their particular roles in shaping the complexity of linguistic features remains a challenge, requiring robust modelling tools to untangle the complex interactions. Here we use an agent-based model to explore for the first time how a combination of prosocial behaviors (here examined from the perspective of ‘self-domestication’), brain development (reflected in the ‘globularization’ of modern skulls) and demographic transitions could have created the context that gave rise to modern linguistic complexity.

The evolution of *Homo sapiens* before the last major out of Africa event around 60 kya is the product of a complex demographic, cultural, environmental and genetic heritage that is not yet fully understood (Scerri et al., 2018). However, the fossil record allows us to identify at least two major transition events that differentiate the anatomy of *Homo sapiens* (Stringer, 2016), relative to their extinct cousins: (i) gracilization/self-domestication: an overall retraction of the face (Lacruz et al., 2019) and (ii) globuralization: a change in brain growth trajectory, leading to an expansion of specific brain regions and giving rise to a more globular cranium (Hublin et al., 2015).

The retraction of the face has been linked to a reduced migration rate and production of neural crest cells (Zanella et al., 2019), features also found in domesticated animals (Wilkins et al., 2014). This major anatomical change has been argued to be the result of a (self-)domestication process (Hare and Woods, 2020), and is accompanied by morphological and behavioral consequences, chiefly a reduction in reactive aggression (Wrangham, 2019) observed across several animal species, such as dogs (*Canis lupus familiaris*), cattle (*Bos taurus*), pigs (*Sus domesticus*), foxes (*Vulpes vulpes*), among others.

It has been independently demonstrated in the Bengalese Finches (*Lonchura striata domestica*), the domesticated variant of the vocal learning White-rumped Munia (*Lonchura striata*) (Okanoya, 2004), that domestication impacts learned behaviors and communicative outputs (see also for data on marmosets reaching a similar conclusion). It is therefore plausible to hypothesize that facial gracilization (self-domestication) enhanced human communicative complexity by boosting prosocial behavior.

The second major anatomical change, globularization, refers to the early period of brain development where the growth of the brain gives rise to the distinctive globular aspect of the skull. This developmental phase, during which “the frontal area becomes taller, the parietal areas bulge, the side walls become parallel, and the occipital area becomes rounder and less overhanging” (Neubauer et al., 2018), is now understood to be the result of a differential growth of brain regions such as the cerebellum, the precuneus and parts of the temporal lobe, and the basal ganglia (Pereira-Pedro et al., 2020). These areas are likely to be associated with modifications of specific memory circuits implicated in handling complex mental sequences, which we find in language (Boeckx, 2017; Gunz et al., 2019; Wynn and Coolidge, 2011).

These two anatomical transitions did not take place at the same time: while the modern face has very ancient roots, with some gracilization traits reaching back to *Homo antecessor* around 1 Mya (Lacruz et al., 2019; Welker et al., 2020; Lacruz et al., 2019), the globularization of the neurocranium appears to be a much more recent event, emerging only in the last 100 kya (Neubauer et al., 2018). For example, the earliest *Homo sapiens* individual, the fossil from Jebel Irhoud (Morocco) dated *ca* 300 kya, has a distinctive modern face but still exhibits an archaic, elongated braincase (Hublin et al., 2017).

Lastly, demography has been repeatedly argued to be critical for understanding the origins of language. In particular, it has been claimed that small populations of speakers exhibit less systematicity (Raviv et al., 2019; Wray and Grace, 2007; Lupyan and Dale, 2010). Similarly, the complexity of material culture has been claimed to be affected by group size (Powell et al., 2009b; Derex et al., 2013; Muthukrishna et al., 2014), with smaller groups struggling to develop and maintain cultural complexity, in part due to drift (but see (Vaesen et al., 2016)).

Here we present a formal model of how these three factors—self-domestication, globularization, and demographic expansion—interact with each other to influence the evolutionary trajectory of linguistic complexity. By modeling the evolutionary mechanisms using simulation, we seek to reveal the importance of the ordering and the contribution of each of these transitions to the evolution of our species’ linguistic capacity. Our approach also allows us to gain insight into the formation of linguistic communities, understood as population clusters with shared vocabularies. Agent-based modelling is a particularly well-suited simulation technique thanks to its ability to integrate individual behaviour leading to emergent outcomes at a population level—such as individual speakers’ interactions culminating in a population level phenomenon: the complexification of language.

## Model description

The model consists of agents characterized by four state variables: `hostility, memorysize, knowns, needs`. At every time step, agents are arranged in pairs at random and individually decide whether or not they wish to communicate with their partner, based on their `hostility`, kinship level, and incentive for communication. Successful communication events lead to transmission of skills and to reproduction, where the child inherits the state variable values of one of the parents chosen at random. Agents live for 30 time steps. A simulation runs for 500 time steps. Output measurements are taken after the last time step has been executed.

Agents are organized in a three-level hierarchical group structure, where every intimate group (5 agents) is a subset of an effective group (25 agents), which is in turn a subset of an extended group (250 agents) (Gamble, 1998). If any group exceeds its maximum size, a new group of the same level is created with half of the agents assigned to it.

At the heart of the model is the agents’ ability to know and exchange *skills.* Skills are represented by integers. Every agent has a list of active skills (`known`) and desirable skills (`needs`) which it wishes (needs) to learn. The known variable is initialized in such a way that two agents share a fraction of their skills proportional to their kinship proximity, i.e. agents’ skillsets within an intimate group will resemble each other more than those within the same extended, but different intimate group. We consider these `known` skills to be an agent’s active vocabulary. The `needs` variable starts empty, increasing by 2 randomly drawn skills at every time step. The maximum length of both the `knowns` and `needs` variables are determined by an agent’s `memorysize`.

For communication to happen, both members of a pair (*A* and *B*) independently calculate their individual cost and incentive for communication.

The *cost* function takes into account the agents’ group relation to its partner as well as its level of hostility. Communication with a distantly related partner (e.g. extended group) is more expensive than with a closely related partner (e.g. intimate group). The extent to which cost increases as a function of group affiliation is proportional to the agent’s hostility. Agents with low hostility will increase their communication cost more moderately than those with high hostility when asked to communicate with a distantly related partner.

The *incentive* function takes into account the percentage of needed skills that the agent can learn from its partner (with a high percentage leading to a high incentive) as well as the number of time steps that have passed since the agent last learned anything (with a longer time increasing the incentive).

The output of the incentive function is subsequently increased or decreased by 2 % depending on whether the partner’s memory size is larger or smaller than the population mean.

If incentive outweighs cost for both agents, then they will communicate. *A* learns all its needs skills that are present in *B*’s `known` variable and vice versa. The length of these transferred signals is one of our key output metrics: *signal length.* A successful communication event produces a child which inherits one of the parents state variable values, chosen at random. In children, the inherited `hostility` and `memorysize` variable values are subject to random mutation, increasing or decreasing by at most 1%. The flow diagram in Fig. 1 provides a visual representation of what happens once two agents have paired up.

**Figure 1:**
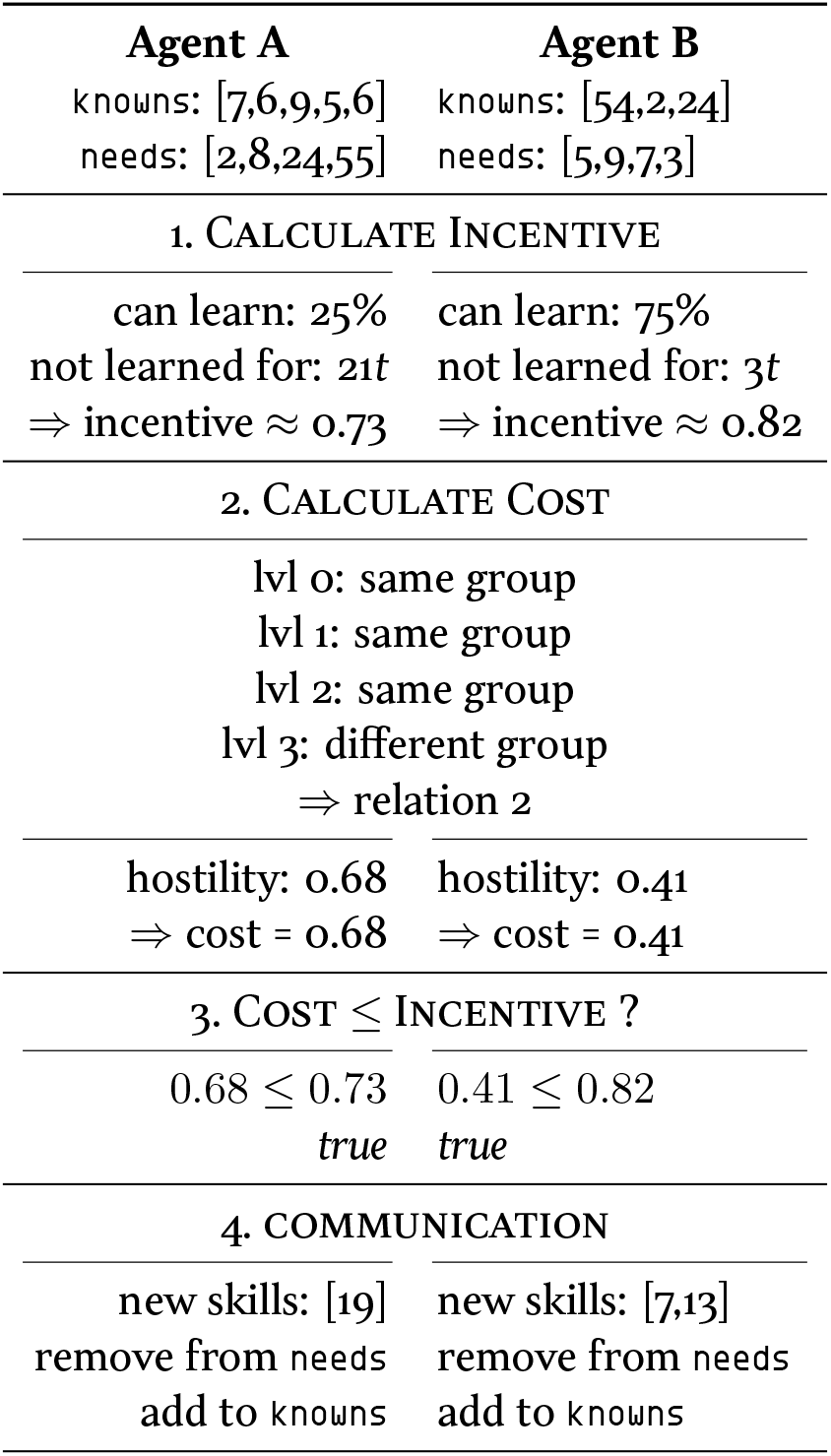
Diagram of communication in the model with example values.

Four global parameters were varied to test a range of scenarios (table 1). `populationsize` is the maximum number of agents that exist at any given time. If the agent population exceeds the parameter value, then the oldest agents are removed until the value is no longer exceeded. `hostility` is the mean of the normal distribution from which agents’ initial hostility is drawn during initialization (std = 0.1). `memorysize` works equivalently (std = 1). `numskills` is the global number of distinct skills that exist in the simulation.

**Table 1:** Tested parameter ranges.

| parameter | default | range | step |
| --- | --- | --- | --- |
| populationsize | 1000 | 100–1000 | 100 |
| hostility | 0.5 | 0.0–1.0 | 0.2 |
| memorysize | 26 | 1–81 | 20 |
| numskills | 26 | 1–81 | 20 |

## Results

Across all runs and parameter configurations, the model reliably showed a significant (*p* < 0.001) decrease in mean `hostility` across the population, at a rate of approximately 0.1 per 500 time steps.

Population growth exceeded death rates under virtually all circumstances. The notable exception, however, was when `memorysize` was low (approx. < 17) causing population growth to stagnate at low values. This is shown in Fig. 2, where population size is very small for low values of `memorysize`, but then suddenly increases, showing a clear exponential relation. With the tested values for `memorysize` being [1, 21, 41, 61, 81] (table 1), only simulations with the lowest value [1] for memorysize were affected by this relation. In all other cases, population growth hit the ceiling imposed by the populationsize parameter. We recognized several recurring patterns across the tested scenarios related to (i) the signal length and (ii) the emergence of linguistic communities.

**Figure 2:**
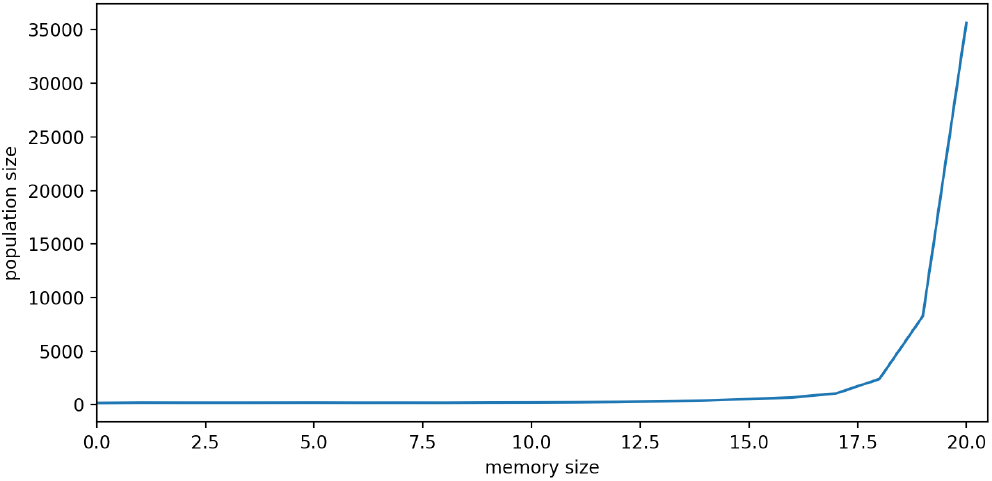
Exponential growth effects of population size in relation to memory size.

### Signal length

An individual signal’s length is defined by the number of skills it contains. One of our key metrics is the mean length of all signals transmitted in the population (measured after the last time step). As a short hand, we refer to this mean value simply as *signal length.* Thus, signal length measures the mean quantity of information shared in individual communication acts across the population. This metric is particularly influenced by both population size and hostility.

Population size influences variation in signal length. As shown in Fig. 3, the variation is high for small populations but, as population size increases, signal length reliably converges until it can almost be considered a constant.

**Figure 3:**
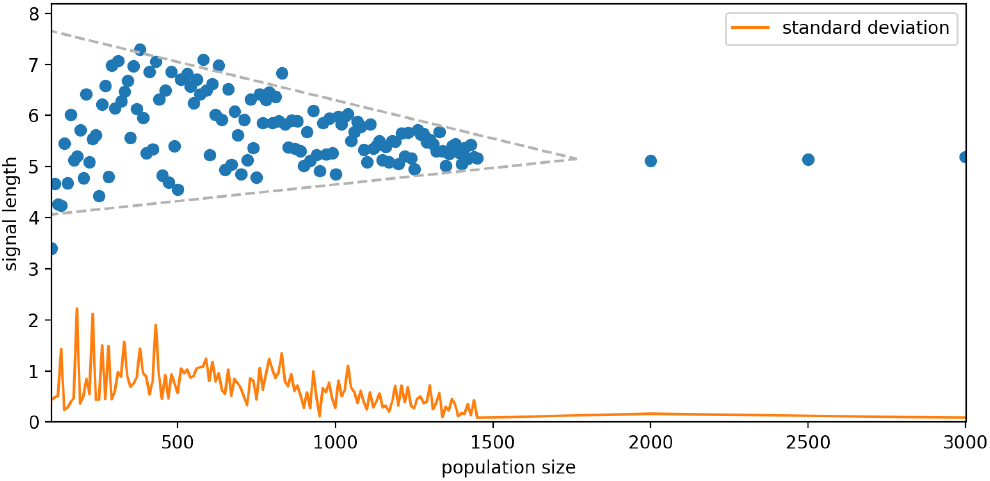
Each data point represents the means of 100 runs with the given population size. Small populations have strong variation in the amount of information they transfer. As population size grows, this variation goes down to virtually zero.

Equally significant is the influence of hostility on signal length, which shows a clear peak of maximum signal length for hostility values around 0.7, above and below which signal length drops (Fig. 4). This result reveals an interesting mechanism arising from the nonlinear interactions between agents.

**Figure 4:**
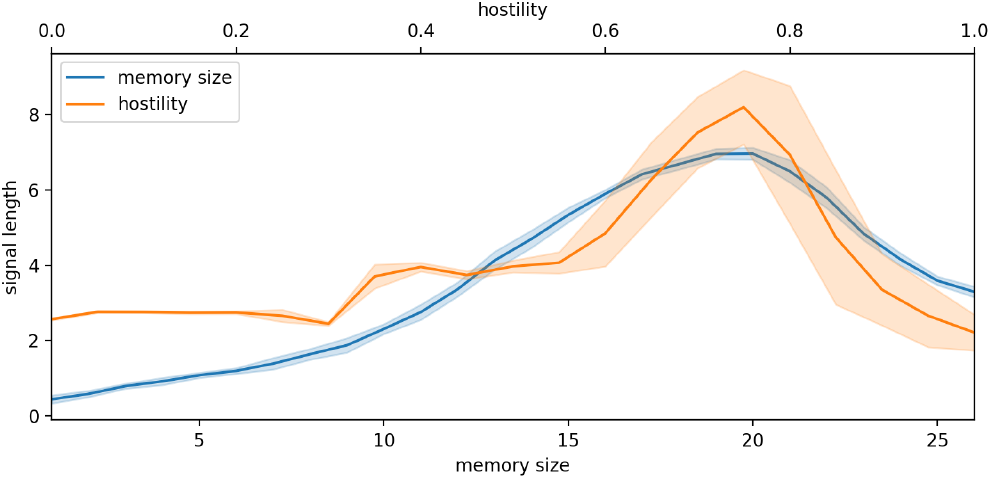
Effects of hostility and memory size on signal length. Both show a clear maximum above and below which signal length decreases.

If hostility is high, the cost of communication outside one’s own intimate group will be high, almost always outweighing the incentive. This leads to communication only happening within, then somewhat isolated, intimate groups. But because agents within the same intimate group also share a majority of their skills, many skills will simply not be present within a given intimate group and communication therefore is limited to short signals. If hostility is low, then in most cases the cost of communication will remain lower than the incentive, leading to frequent communication events. This, in turn, means that agents get to learn all needed skills almost instantaneously and the number of their needs is never large enough to translate into events where long signals can be transmitted.

It must be noted that the position of the maximum, here at 0.75, shifts depending on the values of model parameters. Accordingly, we do not regard the precise position of the maximum to be of great importance, but instead the existence and shape of the maximum itself.

Plotting `memorysize` against signal length shows a curve similar to the one from hostility (Fig. 4). Again, the existence of optimal conditions for `memorysize` can be understood through the interactions of different processes. Below the maximum, `memorysize` acts as an upper bound on the length of agents’ needs, which in turn acts as an upper bound on signal length. Above the maximum, agents will have the majority of skills already stored in their known variable and therefore have no need to learn them, which decreases the amount of skills learned on average and, in turn, decreases signal length.

### Linguistic communities

Our model reveals circumstances where linguistic communities of varying sizes and strengths form. For this analysis, we use the agents’ known variable as a proxy for their individual vocabulary. We applied hierarchical clustering (UPGMA) to the agents, using as our distance metric the edit (Levenshtein) distance between the agents’ (sorted) vocabularies. We define a strongly clustered population (i.e. a population exhibiting linguistic communities) as a population in which there are multiple subsets of agents who share similar vocabulary (low linguistic distance between group members) but the vocabulary in between these subsets differs significantly (high linguistic distance). Such a population will result in a dendrogram similar to Fig. 5b, indicating a clear, distinct jump in linguistic distance between subclusters.

**Figure 5:**
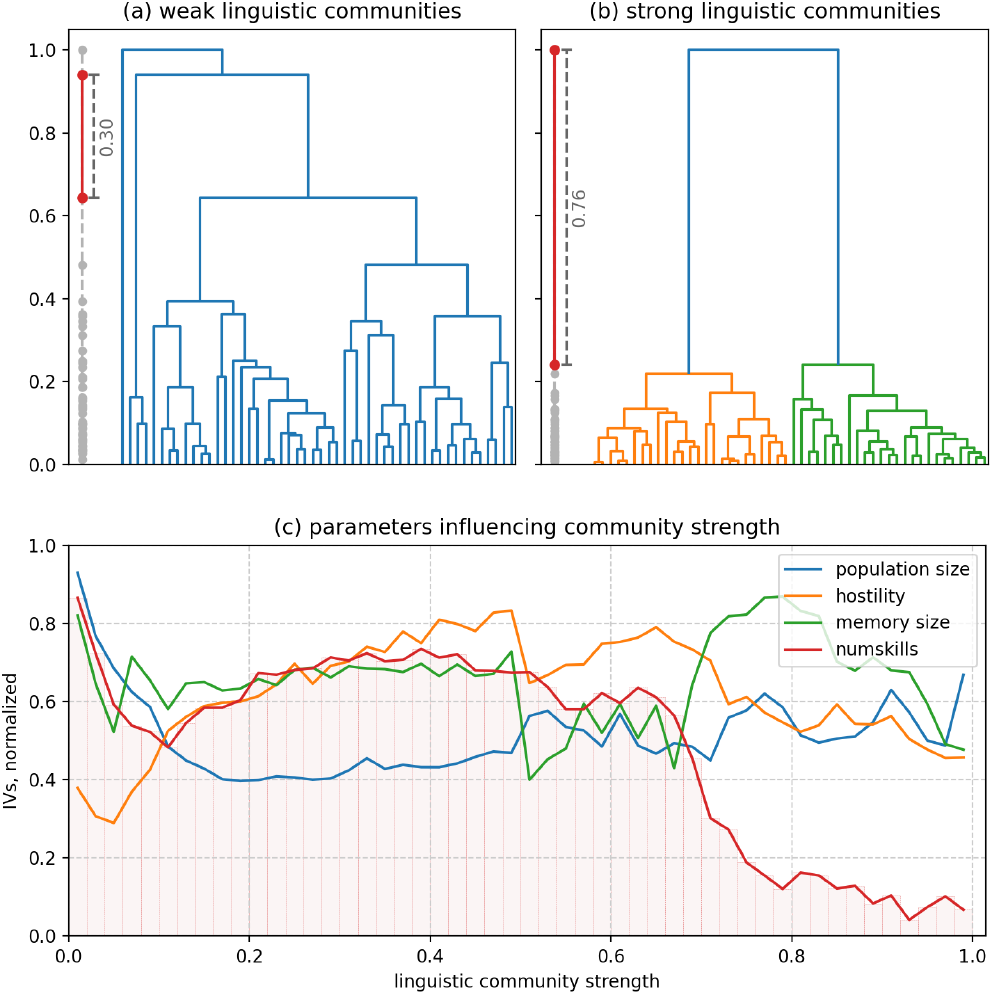
*Top:* We measure the formation of linguistic communities by flattening a normalized dendrogram onto a line and taking the length of the longest segment on this line. Weak linguistic communities have no sudden jumps in vocabulary distance, leading to the longest segment being small; strong linguistic communities do have sudden jumps in vocabulary distance, leading to the longest segment being large. *Bottom:* Parameter values leading to formation of linguistic communities. Data points are bins of size 0.02; all parameters were normalized.

By contrast, we take as weakly clustered a population in which the vocabularies of any two agents have roughly the same edit distance. Such a population will result in a dendrogram similar to Fig. 5a, with gradually increasing distances between subclusters and no sudden jumps. We therefore use the length of the longest segment in a dendrogram, as described in Fig. 5, as a metric indicating how strong the linguistic communities in a population are.

The plot in Fig. 5c plots the strength of linguistic communities against model parameters. The results reveal that linguistic communities are strongest under conditions where the memory size is high but the number of skills is low. In other words, it is under conditions where any individual could easily acquire the entire vocabulary in terms of mental capacity, that we observe the strongest formation of linguistic communities, the strongest competition for vocabulary.

Unsurprisingly, low hostility—which facilitates communication across distant social groups—is related to well mixed populations with no distinct linguistic groups. Increasing hostility translates into stronger linguistic groups only up to a point, after which a further increase shows little impact. This is likely to be directly linked to the same process that causes the global maximum of signal length described before. In contrast, population size has no influence on the strength of linguistic groups.

In addition, we calculated the correlation between an agent pair’s vocabulary similarity and its group similarity across the parameter space. The higher this correlation is, the more does a close group relation coincide with having similar vocabulary. The results of a linear regression analysis, summarized in table 2, show that `memorysize` and `numskills` explain by far the largest part of the variation. In between those two parameters, `numskills` not only has the highest *R*^2^-value, but also a coefficient almost twice as high as that of `memorysize` (0.0043 vs 0.0023). By contrast, both `hostility` and popsize explain little variation, and the latter also shows a lower significance level. These results converge with the hierarchical clustering described previously, indicating that both the formation of linguistic groups as well as their co-occurrence with social groups is chiefly driven by changes in mental capacity at the level of the individual, and much less so by demographic changes at the population level.

**Table 2:** Results of linear regression.

| IV | $F$ | $p$ | $R^2$ |
| --- | --- | --- | --- |
| popsize | $F(1, 734) = 11.61$ | $< 0.001$ | 0.016 |
| hostility | $F(1, 734) = 12.66$ | $< 0.0001$ | 0.017 |
| memorysize | $F(1, 734) = 82.42$ | $< 0.0001$ | 0.101 |
| numskills | $F(1, 734) = 415.3$ | $< 0.0001$ | 0.361 |
| full model | $F(4, 731) = 180.6$ | $< 0.0001$ | 0.497 |

To summarize, the results identify a narrow band of conditions in which the average signal length can increase— a moderate to high level of hostility and moderate range of vocabulary capacity (`memorysize`)—indicating that linguistic expansion could only occur after the processes of self-domestication and globularization were completed. In addition, they suggest a strong effect of memory size on population growth. Finally, the origins of distinctive linguistic communities are also associated with specific circumstances, namely a high vocabulary capacity (`memorysize`) coupled with lower number of individual units present in the language (`numskills`).

## Discussion

Based on the archaeological record and findings in related disciplines researchers have highlighted three important processes in the evolutionary trajectory of our species: self-domestication, globularization and demographic expansion. Each of these three events is incorporated into the model. The agents’ hostility (or lack thereof) serves as a proxy for self-domestication since lower aggression towards strangers is its most characteristic behavioral phenotype (Wrangham, 2019). The increase of agents’ memory size serves as a proxy for globularization (Boeckx, 2017), while an increase in population size is the equivalent to demographic expansion. Through these three proxies, we modelled the underlying processes in as simple as possible terms to investigate the interactions between them and their impact on the evolution of modern linguistic complexity. We do not claim that these proxies capture all the details and aspects of the transitions that took place, but even this simple model reveals how the various factors we modelled conspire to boost linguistic complexity in a highly non-linear and complex way.

All together the model demonstrates the causal chain effect of self-domestication on the evolution of modern linguistic complexity and a temporal ordering of self-domestication causing globularization, which in turn triggers demographic expansion (Fig. 6). A decrease in hostility over time is a robust pattern across all tested scenarios. Decreasing hostility eventually facilitates communication and leads to an increase in signal length. The necessity of higher memory size to transmit longer signals makes it plausible that evolutionary pressure for longer signal transmission manifests itself as pressure for higher memory size and as a result - increased cognitive capacity.

**Figure 6:**
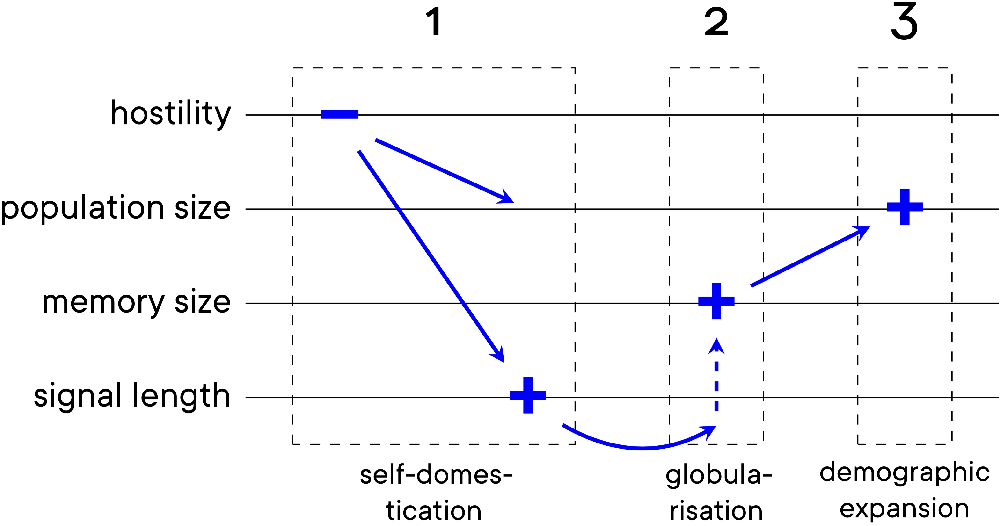
Our model results are compatible with the archaeological record, suggesting a temporal order of the three main transitional events.

As seen in Fig. 2, an increase in memory size correlates with exponential population growth (demographic expansion), completing the sequence of events. Crucially, each parameter in the model was shown to impact distinct aspects of the phenotype, strongly suggesting that a full understanding of linguistic complexity requires taking into account several factors and mechanisms, each impacting the evolutionary process in a specific way and all of them interacting with each other.

The model’s results and in particular the temporal ordering of the processes are highly compatible with the evolutionary timeline that can be inferred from the fossil record. The process of face retraction is well documented already in early hominin lineages, together with an increase in pro-social behaviour - a trend that continues throughout human evolution. Globularization on the other hand is associated with much more recent human past, ca. 350,000-35,000 years ago (Neubauer et al., 2018; Hodgson, 2013). Finally, the end point of the process of endocranial globularisation correlates with the most dramatic demographic expansion of our species approximately 45,000-35,000 years ago (Powell et al., 2009a). The same temporal order of the three processes emerges from our model suggesting that the timing and sequence of these events is not coincidental and, although previously regarded as independent processes, they are in fact closely intertwined.

The emergence of linguistic communities is of wide academic interest. Prior to the use of DNA as the basis of the human phylogenetic tree, linguistic affinity was a primary source of models concerned with the longterm trajectories of human groups (Cavalli-Sforza et al., 1988). In both cases, their association with particular archaeologically or historically identified communities has been a source of much debate (Gimbutas, 1997; Gray and Atkinson, 2003; Haak et al., 2015). The origins of the formation of such distinctive socio-cultural-linguistic units is therefore of great relevance. The diversification into regional “traditions” during the European Upper Palaeolithic has been hailed as one of the landmarks of modern human culture (Gamble, 1998). Our model allows us to gain insight into the conditions that could lead to the development and maintenance of such cultural traditions.

The combination between large language capacity (high `memorysize`) combined with small number of modular language units (`low numskills`) has been suggested as a critical feature of modern human language (Nowak and Krakauer, 1999). Our model indicates that the existence of distinctive language communities constitutes a good proxy for high language complexity.

## Supplementary information

All data, including all code and raw model output data, have been made available under the Creative Commons Attribution-ShareAlike 4.0 license at http://www.doi.org/10.5281/zenodo.4270200.

